# The origin of internal genes contributes to the replication and transmission fitness of H7N9 avian influenza virus

**DOI:** 10.1101/2022.08.17.504359

**Authors:** Joe James, Sushant Bhat, Sarah K. Walsh, H. M. Thusitha. K. Karunarathna, Jean-Remy Sadeyen, Pengxiang Chang, Joshua E. Sealy, Sahar Mahmood, Benjamin C. Mollett, Marek J. Slomka, Sharon M. Brookes, Munir Iqbal

## Abstract

H9N2 avian influenza viruses (AIVs) have donated internal gene segments during the emergence of zoonotic AIVs, including H7N9. We used reverse genetics to generate three reassortant viruses (2:6 H7N9) which contained the Haemagglutinin and Neuraminidase from Anhui/13 (H7N9) and the six internal gene segments from H9N2 AIVs of G1-like or BJ94-like lineages enzootic in different geographic regions in Asia. Infection of chickens with the 2:6 H7N9 containing internal gene segments from G1-like H9N2 conferred attenuation in vivo, with lower shedding and reduced transmission to contact chickens. However, possession of BJ94-like H9N2 internal gene segments resulted in more rapid transmission and significantly elevated cloacal shedding compared to the parental Anhui/13 H7N9. In vitro analysis also showed that the 2:6 H7N9 having BJ94-like internal genes had significantly increased replication compared to the Anhui/13 H7N9 in chicken cells. In vivo co-infection experiments followed, where chickens were co-infected with pairs of Anhui/13 H7N9 and one of each of the three 2:6 H7N9 reassortants. During ensuing transmission events, the Anhui/13 H7N9 virus outcompeted 2:6 H7N9 with internal gene segments of BJ94-like or G1-like H9N2 viruses. Co-infection did lead to the emergence of novel reassortant genotypes that were transmitted to contact chickens. Some of the reassortant viruses had a greater replication in chicken and human cells compared to the progenitors. We demonstrated that the internal gene cassette determines the transmission fitness of H7N9 viruses in chickens and the reassortment events can generate novel H7N9 genotypes with increased virulence in chickens and enhanced zoonotic potential.

**Importance:** H9N2 avian influenza viruses (AIVs) are enzootic in poultry in different geographical regions. The internal genes of these viruses can be exchanged with other zoonotic AIVs, most notably the China-origin H7N9 that can give rise to new virus genotypes with increased veterinary, economic and public health threats to both poultry and humans. We investigated the propensity of the internal genes of H9N2 viruses (G1 or BJ94) in the generation of novel reassortant H7N9 AIVs. We observed that the internal genes of H7N9 which were derivative of BJ94-like H9N2 virus have a fitness advantage compared to those from the G1-like H9N2 viruses for efficient transmission among chickens. We also observed the generation of novel reassortant viruses during chicken transmission which infected and replicated efficiently in human cells. Therefore, such emergent reassortant genotypes may pose an elevated zoonotic threat.

## Introduction

Among the many diverse avian influenza viruses (AIVs), the H9N2 subtype of AIVs became prominent by the turn of the century, where they had become enzootic in poultry in many parts of Asia (1). H9N2 AIVs have continued to evolve in the field, by both genetic drift and by reassortment, whereby the latter frequently shifts the predominant genotypes observed in poultry, with fitter viruses outcompeting existing strains and emerging as novel dominant strains (2, 3). H9N2 viruses can be classified phylogenetically by the haemagglutinin (HA) gene, which include three stable lineages two of which namely G1 and BJ94 (sometimes referred to as Y280), are of Asian origin, the former is essentially restricted to terrestrial poultry but has spread to the Middle East and into Africa (4–6). The third Eurasian grouping, Y439, is part of a broader global lineage which also includes H9N2 AIVs from North American wild birds and European isolates (7, 8). Although all H9N2 AIVs are low pathogenicity AIVs (LPAIVs), they can cause mortality in poultry particularly when associated with secondary microbial infections (9, 10). In addition, H9N2 viruses continue to be responsible for mild zoonotic infections which have been recorded mainly in East Asia since 1998, but with no evidence thus far for any onward efficient transmission among humans (11).

Within the BJ94 lineage, Genotype 57 (G57) H9N2 viruses were first identified in China in 2007, and had become the predominant genotype in poultry due to their enhanced *in vivo* fitness (12). Subsequent reassortment of G57 lineage H9N2 viruses with other circulating subtypes has resulted in the generation of several zoonotic AIVs including: H5N1 (1997-present), H7N9 (2013-present), H10N8 (2014), and H5N6 (2015) (20–23). Some of these novel reassortants contained internal genes from the H9N2 G57 lineage and from several H5Nx highly pathogenic avian influenza viruses (HPAIVs) of the “goose/Guangdong” lineage (13–15). Importantly, it has been previously demonstrated that the internal gene segments of G57 greatly contribute to the pathogenicity of these viruses in mammals, further emphasising their significance as a novel zoonotic threat, whereby their pandemic potential cannot be ignored (13, 14, 16–22).

Following the epizootic emergence of China-origin H7N9 during 2013 (prototype virus strain A/Anhui/1/2013 [H7N9]) through an earlier reassortment involving a wild bird H7 AIV and a G57 H9N2, H7N9 incurred extensively into chickens in China as a LPAIV with an absence of overt clinical signs in chickens (23). This H7N9 virus has caused over 1,500 human infections with a 38% case fatality rate (24), and prompted concerns that it may threaten to evolve into the next human influenza pandemic (25, 26). H7N9 has continued to circulate in poultry and diversified into multiple lineages, in part by dynamically reassorting with other viruses in China, particularly H9N2 viruses (27, 28). Despite widespread H7 AIV vaccination in poultry in China, H7N9 continues to be detected in poultry (24). There is an ever-present concern that geographical expansion of H7N9 could facilitate reassortment with a more diverse range of AIVs, leading to the generation of novel AIVs which are more transmissible or pathogenic in poultry and have a higher propensity for zoonotic transmission to humans (29). Indeed, H9N9 and H7N2 subtypes have already been seen to arise as a result of natural reassortment in the field (30–33) or after experimental co-infection between H7N9 and H9N2 (34), with the progeny having been shown to have increased virulence both experimentally and naturally (34, 35). We have previously demonstrated how a novel H9N9 reassortant virus emerged as the dominant genotype following *in vivo* co-infection between China-origin H7N9 and G1-like H9N2 viruses (34). This H9N9 reassortant virus transmitted successfully to naive chickens, and efficiently replicated in ferrets with evidence of transmission which may have been mediated through aerosols (34). These observations underlined that further reassortment between these H7N9 and H9N2 viruses may result in a significant public health threat, which emphasised the need for further characterisation of such novel reassortments to safeguard both the poultry industry and human health.

We selected different H9N2 viruses representative of the dominant lineages which are circulating in countries neighbouring China. Using reverse genetics (RG) we rescued viruses consisting of the HA and neuraminidase (NA) genes from the prototype China-origin H7N9 (A/Anhui/1/2013, abbreviated to Anhui/13), and the internal gene cassettes from three distinct H9N2 viruses (BJ94-like, G1 subgroup 2 and G1 subgroup 3) isolated from different geographical regions in Asia. We investigated the contribution of the internal genes from H9N2 viruses upon viral fitness of the reassorted H7N9 genotypes in chickens. In order to identify which constellation of H9N2 internal genes may feature in a viable emergent H7N9 reassortant, we also co-infected chickens with mixtures of Anhui/13 and the reassorted H7N9 viruses possessing the different H9N2 internal gene segments. This approach provided elements of competition between the two infecting viruses, along with an experimental setting for the *in vivo* emergence of reassortants. The co-infection experiments prompted characterisation of the viable progeny viruses which included novel genotypes, along with their accompanying consequences for viral fitness which were assessed *in vitro* using chicken cells. Finally, we investigated the potential zoonotic risk of these novel reassortant viruses by characterizing their replication in human lung epithelial cells.

## Results

### The internal gene cassettes of H9N2 viruses from different lineages confer different replication and transmission properties to 2:6 reassortant H7N9 in chickens

We rescued the Anhui/13 (H7N9) virus by reverse genetics (RG). To investigate the effect of other H9N2 internal gene cassettes, we generated three reassortant H7N9 viruses by RG; each virus possessed the surface glycoproteins, HA and NA from Anhui/13 H7N9 and the remaining six ‘internal’ gene cassettes from three genetically different H9N2 viruses. These lineages included two ‘G1-like’ viruses from Pakistan (G1 Subgroup 2-like) and Bangladesh (G1 Subgroup 3-like) and one ‘BJ94-like’ lineage H9N2 virus from Vietnam (Supplementary Figure 1 and Supplementary Figure 2). These viruses are hereby referred to as 2:6 H7N9 viruses (Pakistan_2:6_, Vietnam_2:6_ and Bangladesh_2:6_) (Supplementary Figure 1 B), and were compared for their infectivity, transmissibility and pathogenicity in chickens.

For the single-virus infections, chickens directly infected (D0; “D” denoting “donor”) with the Anhui/13 registered mean positive shedding from the oropharyngeal cavity at 1 day post infection (dpi) with peak virus shedding between 1-4 dpi, which ceased by 8 dpi (Figure 1 A and Supplementary Figure 3). The Vietnam_2:6_, Pakistan_2:6_ and Bangladesh_2:6_ viruses all caused robust infection in D0 chickens (Figure 1 A), producing similar total oropharyngeal shedding profiles to Anhui/13 (Figure 1 E). For all viruses, cloacal shedding was more sporadic than oropharyngeal shedding, observed between 2 and 7 dpi (Figure 1 C). However, cloacal shedding in the D0 Vietnam_2:6_ chickens was significantly greater than chickens infected with Anhui/13 (Figure 1 F). Chickens from all groups exhibited mild-moderate clinical signs with no significant differences observed among the different groups (data not shown).

**Figure 1.**
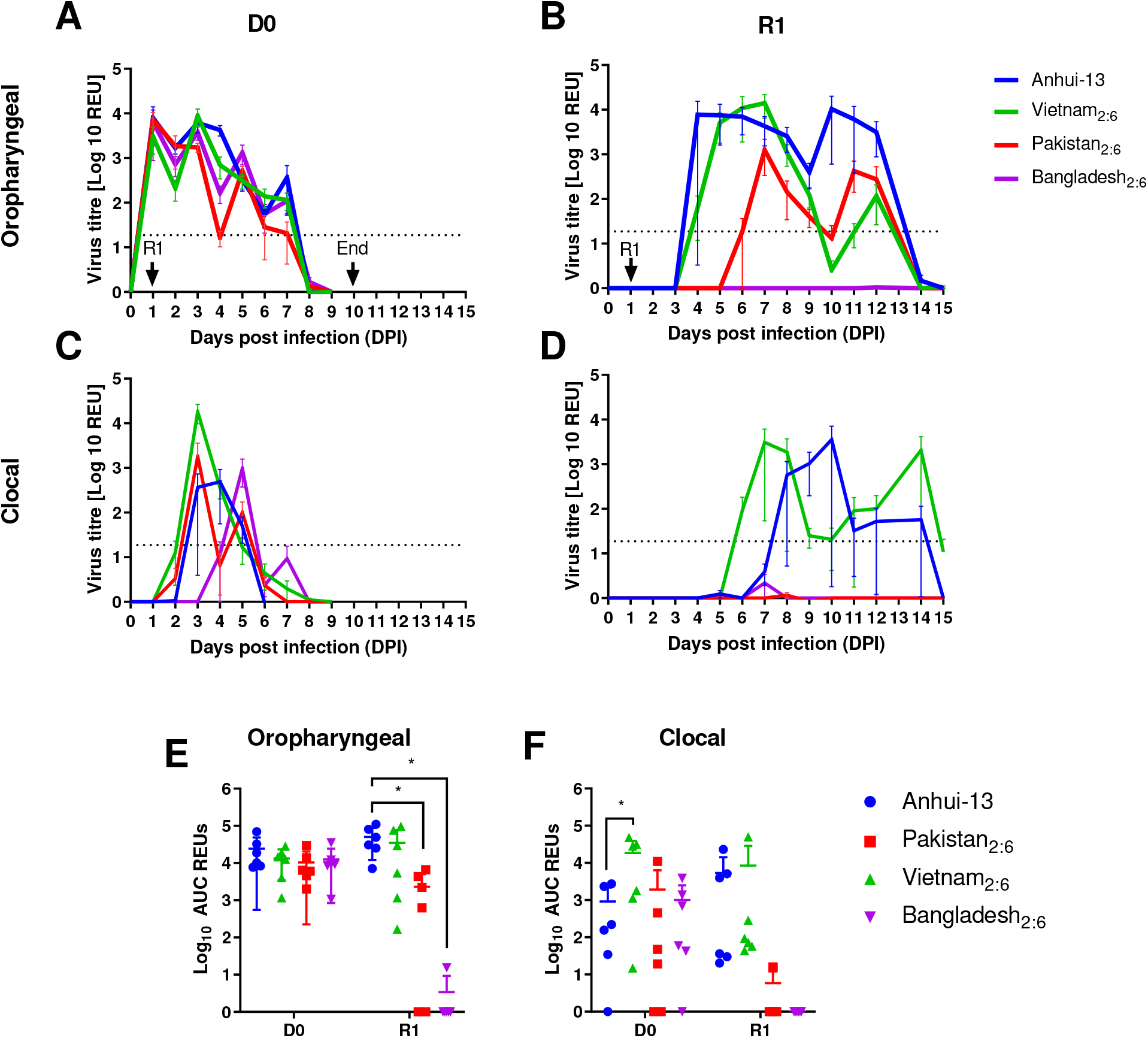
Mean viral shedding and transmission dynamics for Anhui/13 and 2:6 H7N9 viruses possessing different internal genes in chickens. **A & C,** Mean influenza virus titres (shown as Relative Equivalence Units (REUs) of Egg Infectious Dose50 (EID50)) from swabs taken from chickens directly infected (D0) with 1×10^8^ EID_50_ of Anhui (blue) or 2:6 H7N9 viruses possessing the internal genes from either Vietnam (green), Pakistan (red) or Bangladesh (purple) H9N2 viruses. **B & D,** Mean REUs from chickens placed in-contact with infected D0 groups at 1 dpi (R1). Swabs were taken from the oropharyngeal (A & B) or cloacal (C & D) cavities of directly-infected or contact chickens. Each coloured line represents mean shedding +/− SD. Oropharyngeal shedding profiles from individual birds are shown in Supplementary Figure 3. REUs and positive cut-off are described in the Methods section. **E & F**, Area under the curve (AUC) analysis for oropharyngeal (E) and cloacal (F) shedding. Each point represents the AUC for an individual chicken, lines represent mean +/− SD. * indicates p-value <0.05 from a one-way ANOVA with multiple comparisons.

Anhui/13 successfully transmitted to 100% of the naive chickens placed in contact (R1; “R” denoting “recipient”) at 1 dpi, based on virus shedding and seroconversion (Figure 1 B and Supplementary Figure 4). Three R1 chickens shed virus from 4 and 5 dpi with the remaining three shedding from 7 and 8 dpi, after D0 shedding was resolving, suggesting that two rounds of transmission may have occurred (Figure 1 B and Supplementary Figure 3 E). Vietnam_2:6_ also transmitted with 100% efficiency, albeit more rapidly than Anhui/13, where an earlier onset of shedding was apparent (Figure 1 B). The transmission efficiency in the Pakistan_2:6_ group was 66.66% (4 out of 6), with shedding first being observed between 6 and 9 dpi, and 4 out of 6 chickens were seroconverted (Figure 1 B and Supplementary Figure 4). No transmission (0 out of 6) occurred in the Bangladesh_2:6_ group based on shedding and seroconversion (Figure 1 B and Supplementary Figure 4). Only the R1 birds in the Pakistan_2:6_ and Bangladesh_2:6_ virus groups showed a statistically significant decrease in oropharyngeal shedding compared to Anhui/13 (Figure 1 E). Among the R1 birds, only the Anhui/13 and Vietnam_2:6_ groups demonstrated sporadic cloacal shedding (Figure 1D and F), while the other groups did not show any shedding through cloacal route. The Vietnam _2:6._ group showed significantly increased cloacal shedding compared to Anhui/13 group.

Taken together, these results indicated that the internal gene cassettes from the Vietnam H9N2 virus bestow increased cloacal shedding together with more rapid transmission dynamics, along with the same high transmission efficiency observed in the Anhui/13 group. In contrast, the internal gene cassettes from both the Pakistan and, more obviously the Bangladesh H9N2 viruses, resulted in reduced viral shedding and transmission compared to Anhui/13.

### *In vivo* competition experiments in chickens co-infected with a mixture of Anhui/13 H7N9 and the 2:6 reassorted H7N9 viruses

We next performed a series of *in vivo* co-infection experiments using Anhui/13 H7N9 and each of the 2:6 H7N9 viruses (Pakistan_2:6_, Vietnam_2:6_ or Bangladesh_2:6_) which were administered to D0 chickens. Two subsequent rounds of contact chickens (R1 and R2) were introduced in an attempt to establish an onward chain of transmission. D0 chickens co-infected with the Anhui/13 and each 2:6 H7N9 virus demonstrated a similar oropharyngeal and cloacal shedding pattern compared to Anhui/13 alone (Figure 2 A-D). The R1 chickens introduced at 1 dpi, began shedding from the oropharyngeal cavity from 3 dpi (2 dpc) (Figure 2 E-H). Based on viral shedding and seroconversion, transmission efficiency to the R1 chickens was 100% in all groups (Figure 2 E-H and Supplementary Figure 5). A group of second contacts (R2) were added upon removal of the D0 chickens. Again, viral shedding indicated 100% transmission efficiency to the R2 chickens in all groups (Figure 2 I-L). However, not all the R2 chickens seroconverted to H7N9 antigen (Supplementary Figure 5), likely because blood sampling was carried out at 14 dpi and some R2 chickens couldn’t seroconvert early. Cloacal shedding was detectable, yet sporadic, for all groups and typically mimicked oropharyngeal shedding, typically commencing at 2 days after onset of oropharyngeal shedding (Figure 2 M-X).

**Figure 2.**
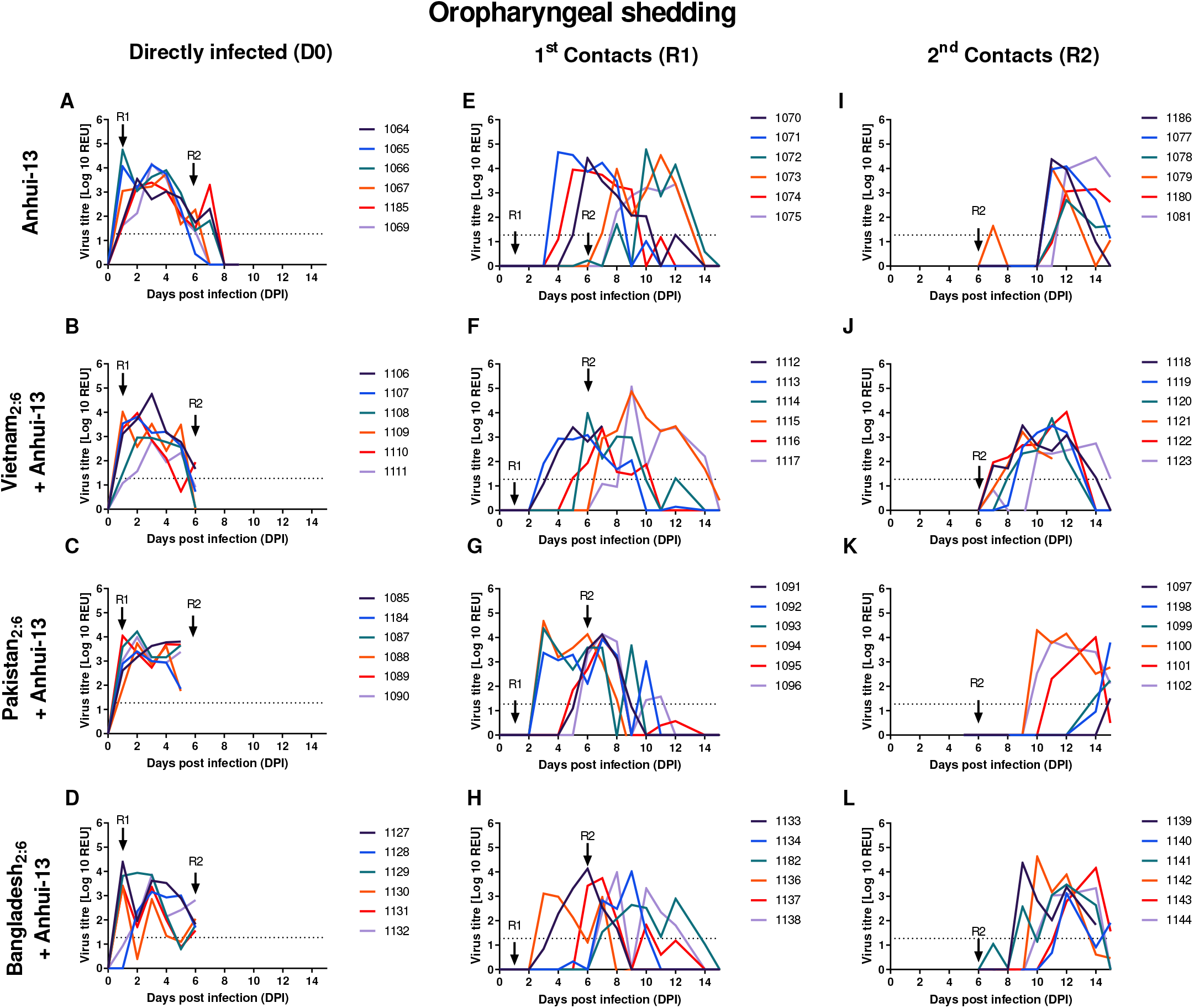
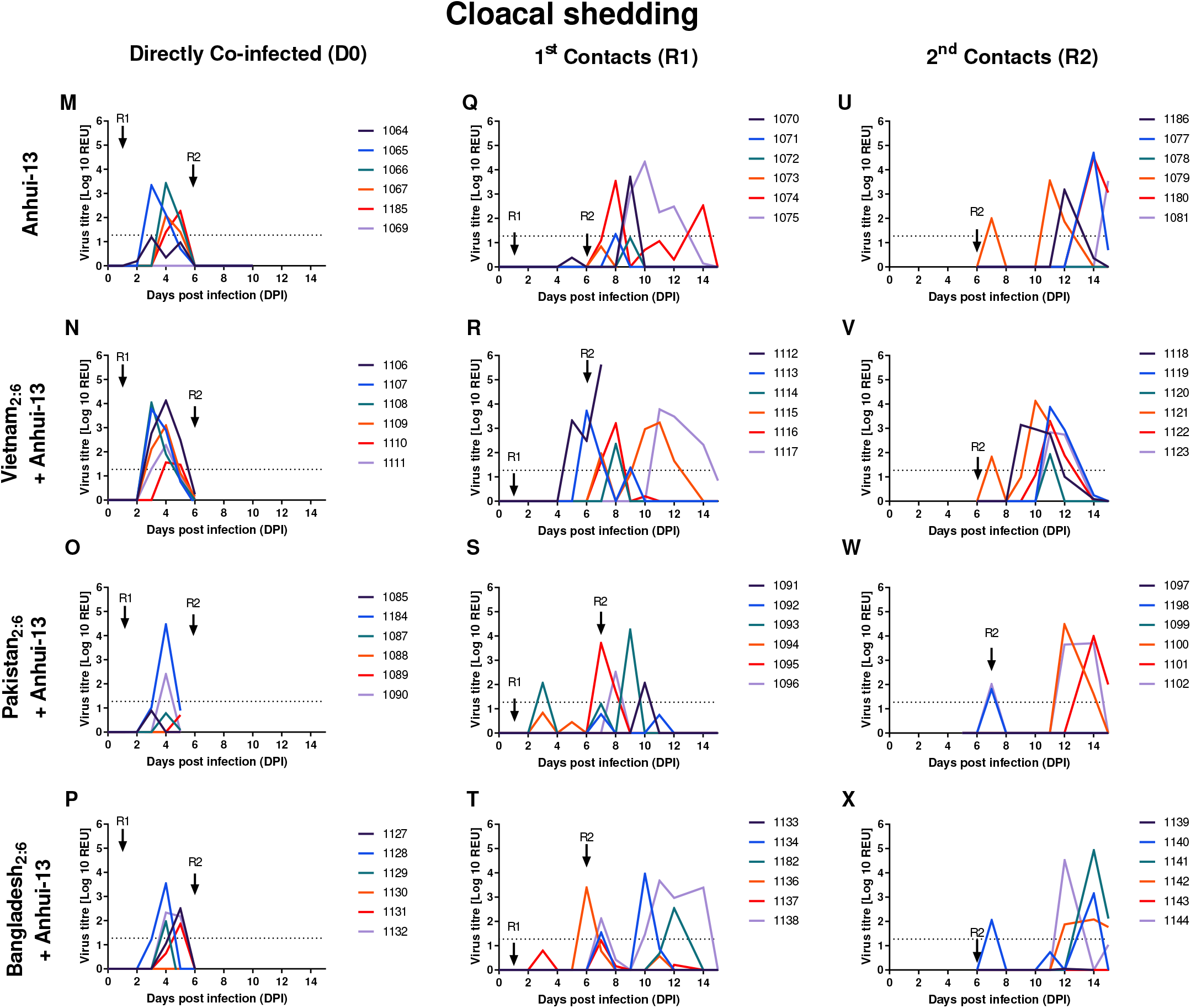
Viral shedding from individual chickens co-infected with both Anhui/13 and one of the three 2:6 H7N9 viruses possessing different internal genes and transmission to contact chickens. Influenza virus titres (shown as Relative Equivalence Units (REUs) of Egg Infectious Dose50 (EID50)) from (A-L) oropharyngeal or (M-X) cloacal swabs from chickens infected with 1×10^8^ EID_50_ of Anhui (A, E, I, M, Q, U) or co-infected with 1×10^8^ EID_50_ of Anhui and 1×10^8^ EID_50_ of 2:6 H7N9 viruses possessing the internal genes from either Vietnam (B, F, J, N, R, V), Pakistan (C, G, K, O, S, W), or Bangladesh (D, H, L, P, T, X) H9N2 viruses. Shedding titres from chickens placed in-contact with infected groups at 1 dpi (1^st^ contacts [R1]) (E-H, Q-T; oropharyngeal and cloacal respectively) or placed in-contact with the 1^st^ contacts (2^nd^ contacts [R2]) (I-L, U-X; oropharyngeal and cloacal respectively). Each coloured line represents shedding from an individual chicken. Arrows indicate the times at which R1 and R2 contact chickens were introduced for successive cohousing periods. REUs and positive cut-off are described in the Methods section.

To investigate the constellation of viral gene segments in the R1 and R2 groups following initial D0 co-infection, lineage specific RT-qPCRs were developed for each internal gene segment (PB2, PB1, PA, NP, M and NS). This approach enabled the specific quantification of each of the segments from Anhui/13 and the six internal genes of the three 2:6 H7N9 viruses.

The viruses used for the inocula displayed equal (~50%) abundance of Anhui/13 and each 2:6 H7N9 virus gene segment (Figure 3). In oropharyngeal swabs from D0 chickens at 1 dpi, both Anhui/13 and 2:6 H7N9 internal gene segments were identified in all groups, albeit at different relative abundances (Figure 3). Interestingly, oropharyngeal swabs sampled at 1 dpi from the ‘D0’ chickens from the Anhui/13 + Vietnam_2:6_ (Figure 3A) and Anhui/13 + Bangladesh_2:6_ (Figure 3C) groups displayed an increased presence of Vietnam- and Bangladesh-origin internal genes, respectively. However, a decline in the frequency of H9N2 internal gene segments was already apparent within these oropharyngeal swabs at the later (4 dpi) D0 shedding stage. Indeed, Anhui/13 origin gene segments were thereafter predominantly detected at the R1 stage, and by the R2 stage appeared to have successfully out-competed the other H9N2-origin segments, regardless of the mixture administered at the initial D0 co-infection (Figure 3). The restoration of what appeared to be a pure Anhui/13 genotype at the R2 stage following all three initial D0 co-infections was observed in both oropharyngeal and cloacal swabs. It was clear that onward *in vivo* transmission in chickens had affected a powerful selection which favoured the Anhui/13 internal gene cassette.

**Figure 3.**
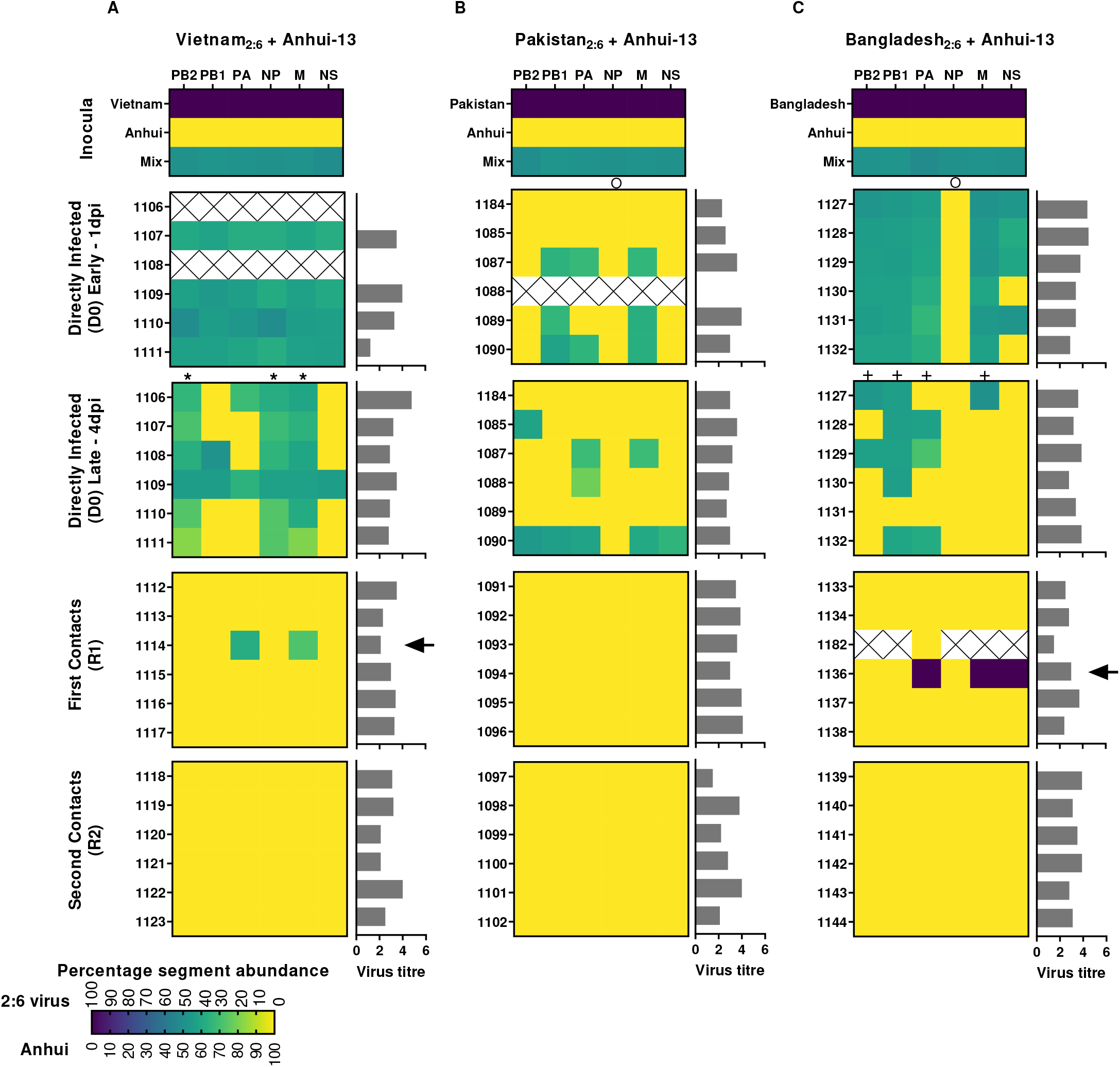
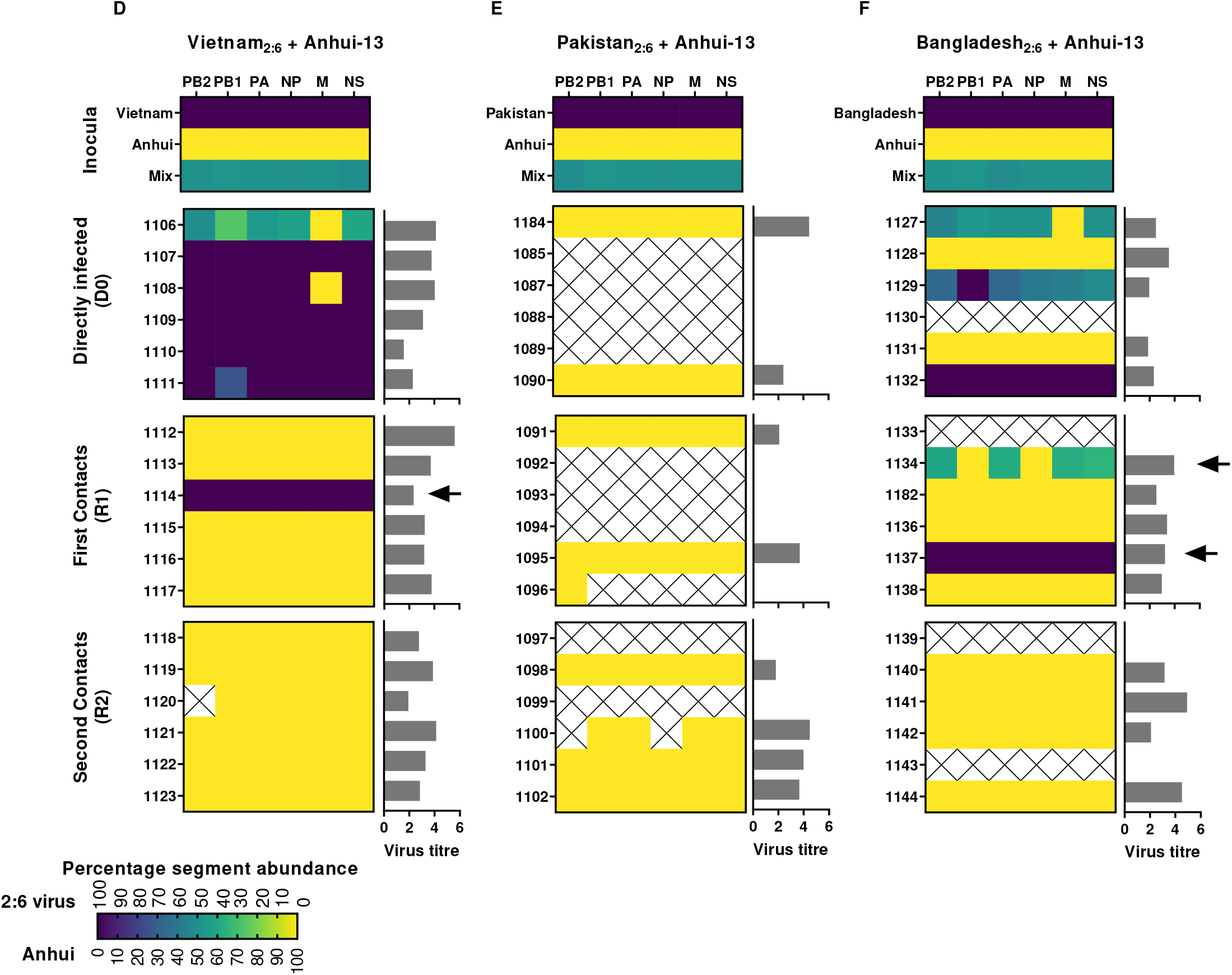
Proportions of distinct internal gene segments among potential progeny viruses from chickens co-infected with Anhui/13 and three 2:6 H7N9 viruses possessing different internal genes. Progeny viruses were identified at the directly infected (D0, divided into early [1dpi] and late [4dpi] for the oropharyngeal shedding) and at the first (R1) or second (R2) contact stages. Swabs from the highest peak viral shedding were analysed unless stated. Heat map colours denote the relative percentage abundance of each internal gene segment from viruses in oropharyngeal (A-C) and cloacal (D-F) swabs obtained from the three consecutive transmission stages. 100% Anhui/13 segment abundance is shown in yellow; 100% 2:6 virus gene abundance is shown in purple; mixed gene abundance is denoted by blended colours. Crosses indicate no gene detection for either virus. Quantified viral shedding titres for each of the analysed swabs are indicated by the horizontal grey bars (REUs, as determined by the M-gene RRT-qPCR). Arrows, asterisks, circles and plus symbols indicate gene segments of interest noted in the Results section.

### Detection of other potential reassortants at the D0 and R1 stages of transmission

Prior to the exclusive emergence of Anhui/13 at the R2 stage in all three groups, internal segments from all three 2:6 H7N9 viruses were detected during the D0 and R1 stages at different frequencies. These observations suggested that several potential reassortant viruses emerged as part of a transient mixed population of progeny viruses (Figure 3). In the Vietnam_2:6_ + Anhui/13 co-infected group, mixed PB2, NP and M gene segments were detected in the oropharyngeal samples at the later D0 stage (Figure 3A – asterisk). At the R1 stage, one potential reassortant was identified from chicken 1114 which contained the PA and M segments from Vietnam at a relative percentage abundance of 38.5 and 28.5% respectively, with the remaining six gene segments from Anhui/13 at 100% abundance (Figure 3A – arrow). Interestingly, the same chicken also shed virus consisting of purely Vietnam internal segments from the cloacal cavity (Figure 3D).

In the Pakistan_2:6_ + Anhui/13 co-infected group, potential reassortment was lower that for the other two groups, where this difference was discernible during oropharyngeal shedding at the early D0 stage (compare Fig 3B with Fig 3A and C). Pakistan_2:6_ internal segments were not detectable at any of the transmission stages (R1 or R2) from both shedding tracts (Figure 3B and E). While chicken #1090 showed potential reassortant viruses in all gene segments, except NP, at the later D0 stage (4dpi), three other chickens in this group also showed generation of single or double potential gene reassortants in the PB2, PA or M gene segments. However, the cloacal samples showed either only Anhui/13 H7N9 origin gene segments or, due to low/undetectable virus titres, genotyping could not be performed.

The oropharyngeal samples at the late D0 stage in the Bangladesh_2:6_ + Anhui/13 co-infected group displayed potential reassortant viruses containing PB2, PB1, PA and M segments originating from Bangladesh_2:6_ with varying abundance (Figure 3C – plus). The cloacal samples from some ‘D0’ chickens included shedding of internal genes of Bangladesh_2:6_ (chicken 1137) or Anhui/13 origin (chicken 1128 and chicken 1131) with 100% abundance (Fig 3F). The other D0 chickens predominantly shed either a mix of internal genes from Bangladesh_2:6_, or due to the low virus titer, the genotype could not be identified. The R1 stage oropharyngeal samples revealed one potential reassortant in chicken 1136 which contained the PA, M and NS gene segments from Bangladesh (with a 100% abundance for all three segments) and all other gene segments from Anhui/13 H7N9 (with 100% abundance) (Figure 3C – arrow). Cloacal swabs at the R1 stage revealed Bangladesh internal segments in two chickens (1134 and 1137) (Figure 3F – arrow). Chicken 1134 included the PB2 (59.3%), PA (61.5%), M (62.2%) and NS (65.1%) gene segments of Bangladesh-origin, with the remaining gene segments from Anhui/13. Chicken 1137 contained all six internal genes from Bangladesh with 100% abundance (Figure 3F). The potential viral progeny produced in the respiratory and enteric tracts of R1 chickens 1134 and 1137 clearly differed as regards the detected frequency of the various gene segments.

In the Pakistan_2:6_ + Anhui/13 and Bangladesh_2:6_ + Anhui/13 co-infected groups, the potential reassortants identified in the oropharyngeal samples of all D0 chickens included the NP gene originating from Anhui/13 H7N9, which was maintained throughout subsequent transmission (Figure 3B and C – circles). The D0 cloacal samples from the Bangladesh_2:6_ + Anhui/13 co-infected group, however, did include shedding of NP from Bangladesh_2:6,_ but again any potential reassortants were outcompeted by viruses containing internal genes of Anhui/13 H7N9 origin during the ensuing transmission stages (Figure 3F). These results showed that NP originating from any other H9N2 virus was rapidly outcompeted by the Anhui/13 NP in both the respiratory and intestinal tract.

### Plaque-purification of viable reassortant progeny viruses from contact chickens

To confirm the exact gene constellation of any viable reassortant viruses which successfully transmitted to contact chickens, we performed plaque purification on swabs from R1 chickens which contained detectable 2:6 H7N9 virus gene segments. The selected specimens included oropharyngeal swabs from chickens 1114 and 1136, plus cloacal swabs from chickens 1114, 1134 and 1137 [Figure 3 – arrows]). We isolated at least 24 plaques per swab and used the segment specific RT-qPCRs to determine the genetic constellation of each isolated plaque. The plaque purification findings matched the RT-qPCR genotyping of the corresponding swab specimens, and therefore confirmed the viability of *bona fide* reassortant viruses (Figure 3 and Table 1). In total, five novel reassortant viruses, with different genetic constellations, were recovered from swabs collected at the R1 stage (H7N9-R1-V29, 30, 31, 32 and 33). The Bangladesh_2:6_ co-infected group produced three new genotypes, one virus as the only population in the oropharyngeal swab from chicken 1136 (genotype H7N9-R1-V29) which contained the Bangladesh-origin PA, M and NS with the remaining gene segments from Anhui/13 (Table 1). Variations of this genotype were also shed from the cloacal cavity of chicken 1134, including viruses containing the Bangladesh-origin PB2, PA, M and NS (genotype H7N9-R1-V33), and PB2, PA and M (genotype H7N9-R1-V30), with the remaining gene segments from Anhui/13. The Vietnam_2:6_ co-infected group produced two new genotypes from the oropharyngeal cavity of chicken 1114, these viruses contained the Vietnam-origin PA (genotype H7N9-R1-V32) plus PA and M genes (genotype H7N9-R1-V31), with the remaining gene segments from Anhui/13 (Table 1). Unaltered Vietnam_2:6_ and Bangladesh_2:6_ (together with unaltered Anhui/13) were also identified by plaque purification, which again reflected the possible viable genotypes indicated earlier (Figure 3 and Table 1).

**Table 1.**
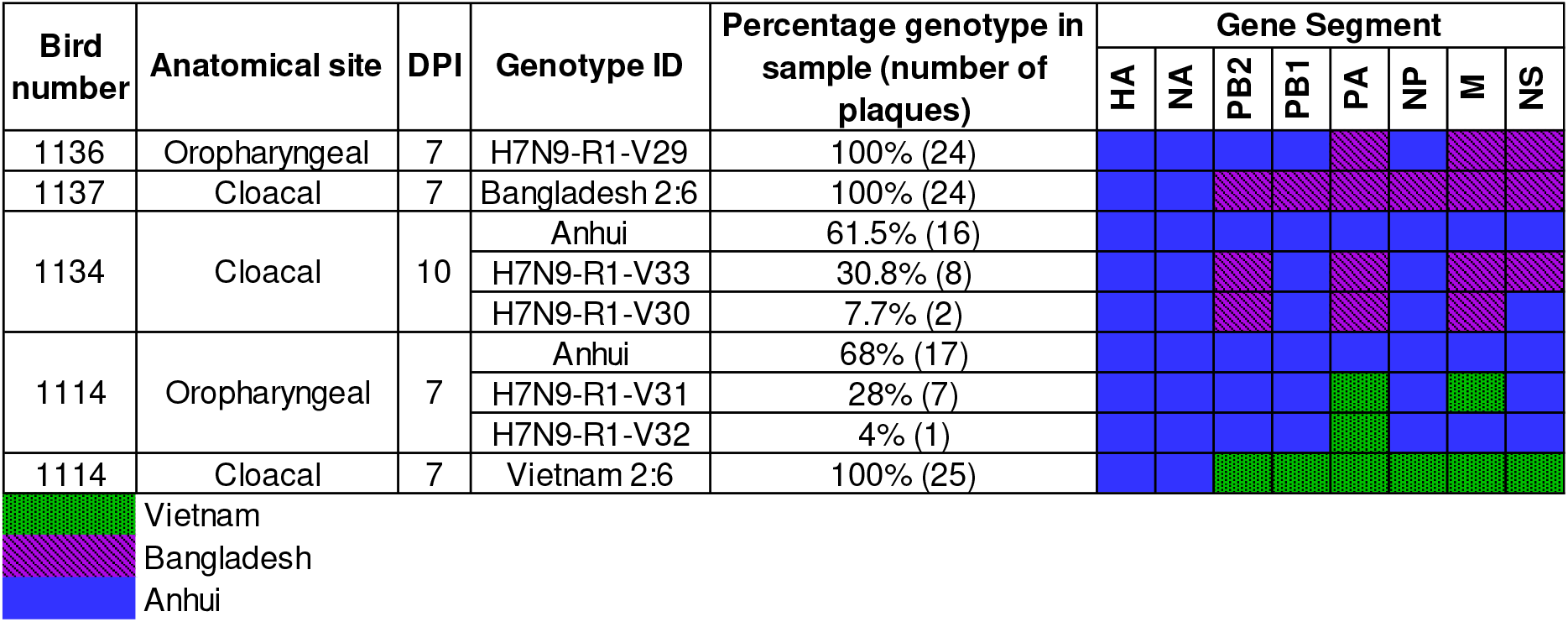
The genetic constellation via plaque purification of potential reassortant viruses identified via RT-qPCR from swabs of R1 contact birds

The Anhui/13 internal genes featured as the exclusive internal gene cassette in all plaque-recovered viruses at the R2 stage (data not shown). However, the R1 transmission stage included plaque-recovered viruses possessing genetically different internal genes as viable novel reassortants.

### Replication of reassortant viruses in chicken and human cells

In addition to the three 2:6 H7N9 viruses which featured in the chicken co-infections, RG was also used to generate the five novel reassorted genotypes which emerged at the R1 stage following co-infection, namely H7N9-R1-V29-H7N9-R1-V30, H7N9-R1-V31, H7N9-R1-V32 and H7N9-R1-V33 (Table 1). In chicken cells (DF-1) there was no statistically difference in replication kinetics between Bangladesh_2:6_ and Pakistan_2:6_ compared to Anhui/13 H7N9 (Figure 4A and C). The Vietnam_2:6_ virus showed significantly higher replication rate compared to Anhui/13 at 24 hrs (P<0.005) and 72 hrs (P<0.001) post-infection (Figure 4B). Among the five reassortant genotypes only two showed statistically different replication kinetics compared to Anhui/13 H7N9. R1-V30 (Bangladesh PB2, PA and M) showed reduced replication in chicken cells at 48 hrs post-infection (P<0.005) (Figure 4E). However, R1-V31 (Vietnam PA and M) showed significantly higher replication rate compared to Anhui/13 H7N9 (P<0.005 at 24 and 48 hrs; P<0.001 at 72 hrs). Further, H7N9-R1-V32 (Vietnam PA) showed comparable replication to Anhui/13 H7N9 indicating that M in combination with PA increase the replication of H7N9-R1-V31. These results suggest that internal gene segments from G1 lineage H9N2 viruses have no effect on the replication of 2:6 H7N9 viruses in chicken cells. However, some combination of G1 gene segments may reduce the replication of 2:6 H7N9 compared to Anhui/13 H7N9 virus. The internal gene segments from the Vietnam-origin (G57 lineage) H9N2 increased the replication rate of 2:6 H7N9 compared to Anhui/13 H7N9 virus, which could be attributed at least due to M gene segment.

**Figure 4.**
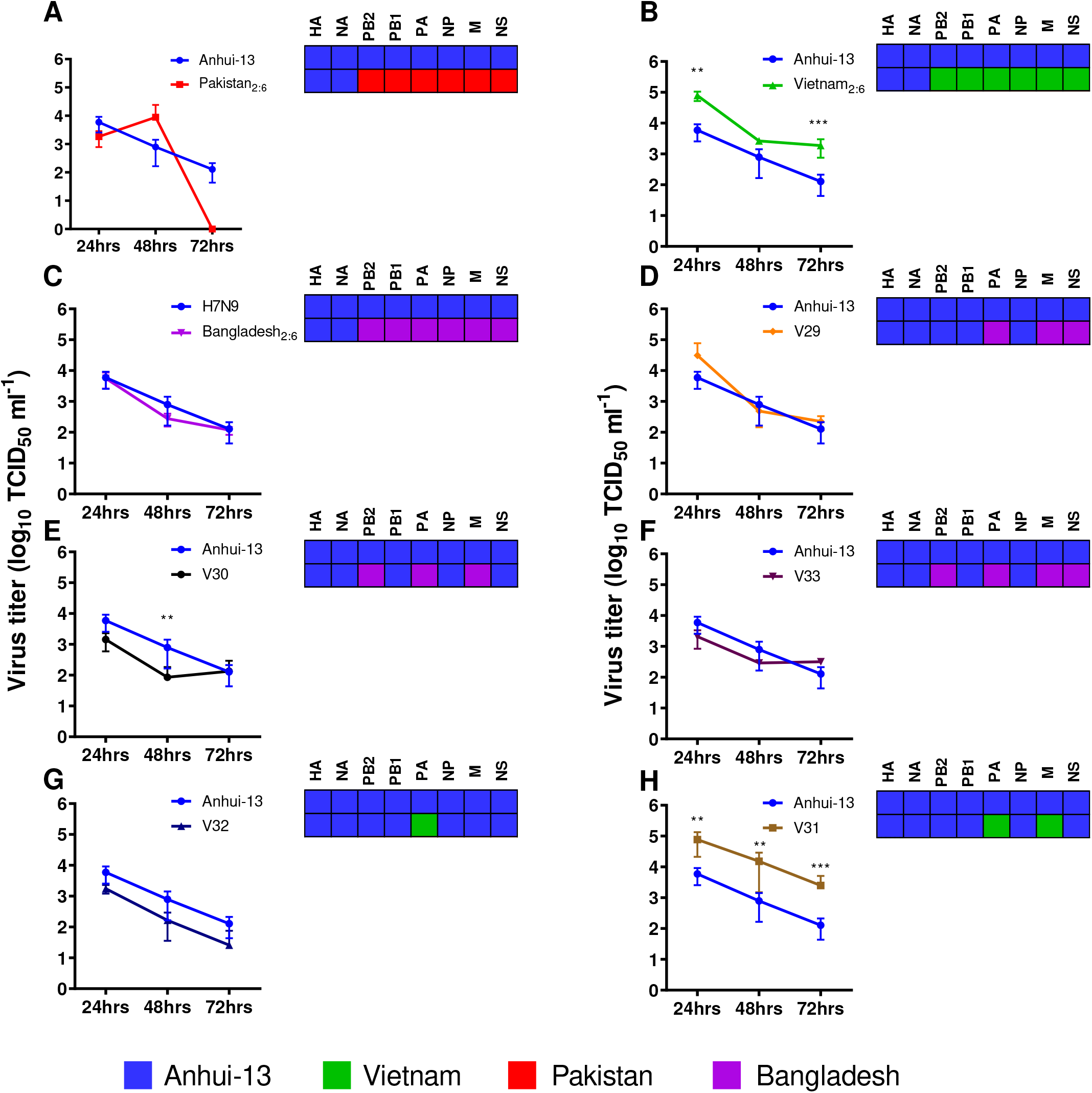
Multi-step replication kinetics of Anhui/13 H7N9, 2:6 H7N9 viruses and the reassortant viruses in chicken DF-1 cells. Virus replication of Anhui/13 (blue) compared to 2:6 H7N9 viruses containing the HA and NA from H7N9 and the remaining internal gene cassettes from either Pakistan (A), Vietnam (B) or Bangladesh (C) H9N2 viruses or novel reassortant viruses recovered by plaque purification from R1 chicken swabs (D-H). Cells were infected at a MOI of 0.01 with each virus and cell supernatant was harvested at 24, 48 and 72 hours post infection. Viral titres were determined by TCID_50_. Each time point corresponds to the mean of four biological replicates with standard deviations indicated. Replication kinetics of reassortant viruses compared to parental Anhui/13 H7N9 virus is shown in A to H. The genotype of each reassortant virus is shown as a combination of black, red, green and violet colours. Two-way ANOVA with multiple analysis was performed comparing every group to Anhui; ** indicates P-value < 0.005, *** indicates P-value ≤ 0.001.

In human cells (A549) Anhui/13 replicated to similar titres compared to Pakistan_2:6_, Vietnam_2:6_ and two reassortant genotypes (H7N9-R1-V31 and H7N9-R1-V32) both containing Vietnam_2:6_ internal gene segments (Figure 5A, B, G and H). However, one reassortant virus, H7N9-R1-V29 (Bangladesh PA, M and NS), had significantly elevated replication kinetics at all timepoints compared to Anhui/13 (Figure 5D). All other viruses replicated to significantly lower titres for at least one time-point compared to Anhui/13 (Figure 5D). In another human cell type (Calu-3) only H7N9-R1-V29 showed comparable replication to Anhui/13 H7N9 (Supplementary Figure 8). Together this suggests that H7N9 virus acquiring ‘all’ internal gene segments from G1 lineage or G57 lineage of H9N2 viruses via reassortment may potentially compromise its replication competence in human cells or the replication rate may remain unaffected. Thus, reassortant 2:6 H7N9 (Pakistan_2:6_, Vietnam_2:6_, and Bangladesh_2:6_) may carry either reduced or similar zoonotic potential as Anhui/13 H7N9. However, some reassortants (possessing PA, M and NS from Bangladesh_2:6_) exhibited increased or comparable replicative fitness in human cells (in Calu-3 or A549) compared to Anhui/13 H7N9, and therefore may represent a potential zoonotic threat.

**Figure 5.**
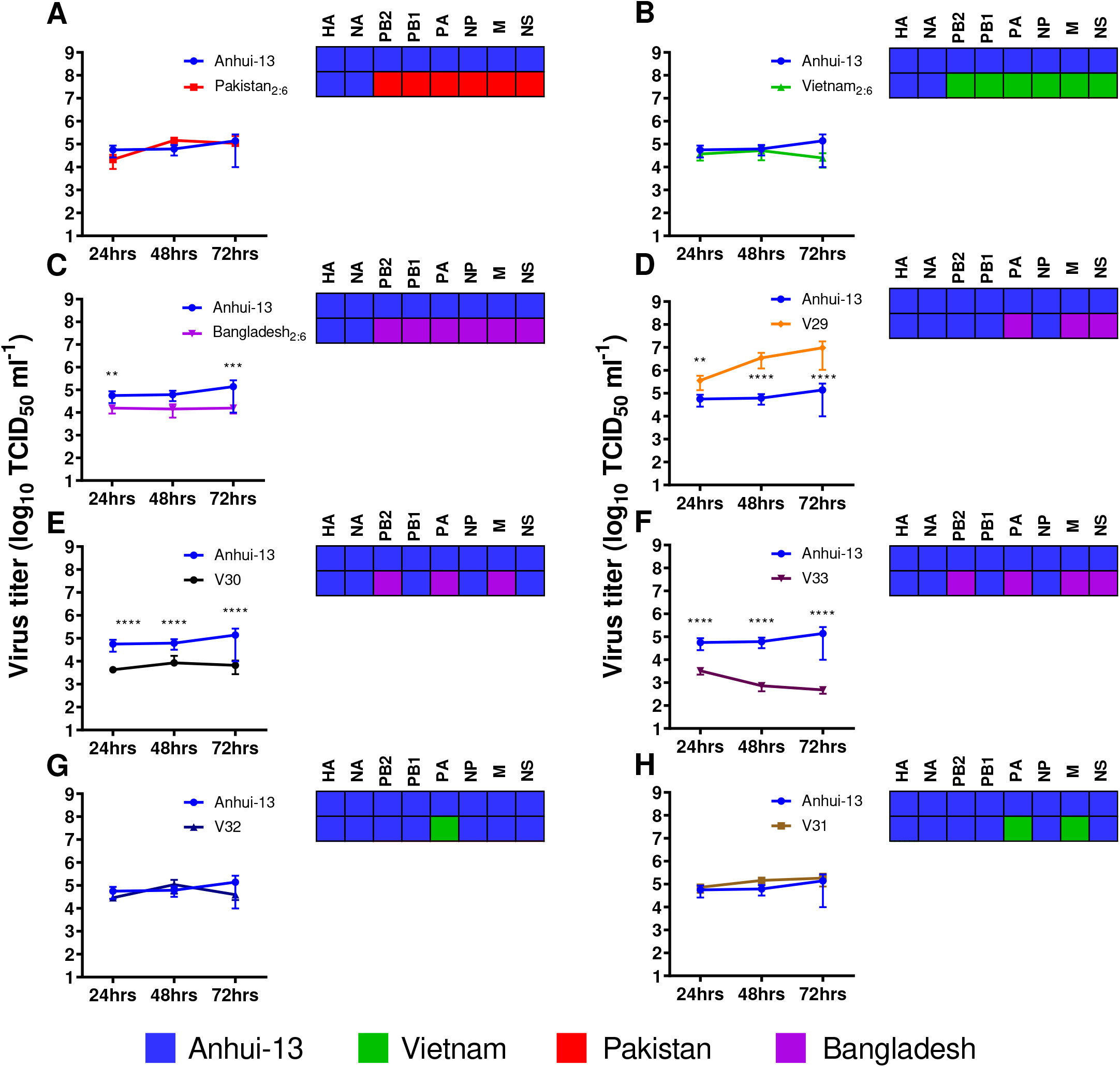
Multi-step replication kinetics of Anhui/13 H7N9, 2:6 H7N9 viruses and the reassortant viruses in human A549 cells. Virus replication of Anhui/13 (blue) compared to 2:6 H7N9 viruses containing the HA and NA from H7N9 and the remaining internal gene cassettes from either Pakistan (A), Vietnam (B) or Bangladesh (C) H9N2 viruses or novel reassortant viruses recovered by plaque purification from R1 chicken swabs (D-H). Cells were infected at a MOI of 0.01 with each virus and cell supernatant was harvested at 24, 48 and 72 hours post infection. Viral titres were determined by TCID_50_. Each time point corresponds to the mean of four biological replicates with standard deviations indicated. Replication kinetics of reassortant viruses compared to parental Anhui/13 H7N9 virus is shown in A to H. The genotype of each reassortant virus is shown as a combination of black, red, green and violet colours. Two-way ANOVA with multiple analysis was performed comparing every group to Anhui; * indicates P-value = 0.0113; ** indicates P-value = 0.0012, **** indicates P-value <0.0001.

### Polymerase activity of reassortant viruses in chicken and human cells

The viral RNA-dependent-RNA polymerase (vRNP) complexes of both Vietnam and Pakistan origins showed significantly higher polymerase activity compared to that of Anhui/13 in chicken cells (DF-1) (Figure 6A). Comparatively, the vRNP complex of Bangladesh showed very low polymerase activity. We also compared the polymerase activity of reassortant viruses identified after plaque purification. The PA alone, or in combination with PB2 from Bangladesh and other RNPs from Anhui/13, also showed significantly lower polymerase activity in chicken cells (Figure 6A). This observation suggested that the Pakistan and Vietnam RNP complexes are more active, while the Bangladesh RNP complex is relatively less dynamic in avian cells.

**Figure 6.**
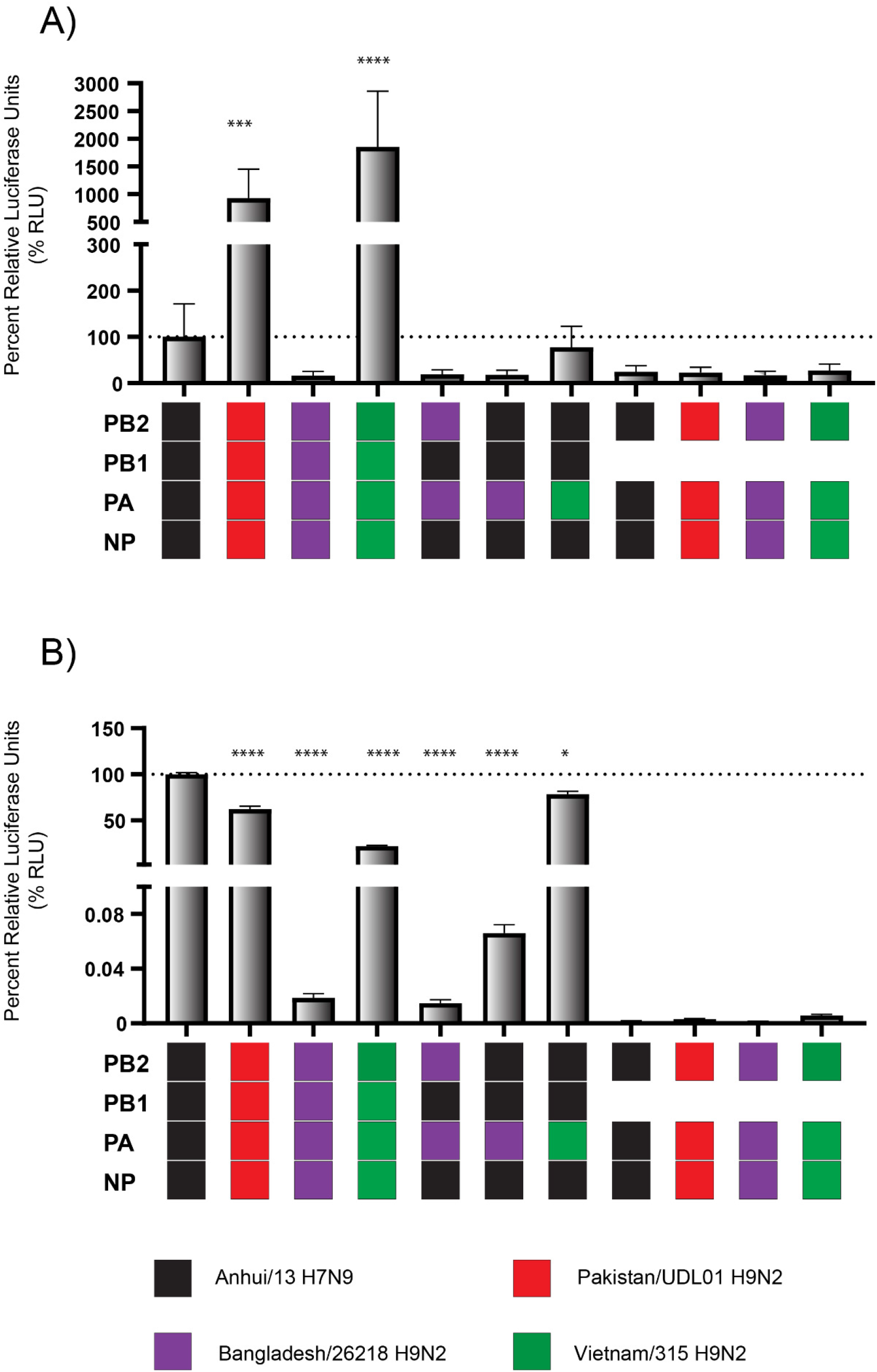
Minireplicon assays of the ribonucleoprotein (RNP) complexes of H7N9 Anhui/13 and H9N2 UDL/08, H9N2 Vietnam/315, H9N2 Bangladesh/26218 and three altered RNP combinations. The RNP complexes were reconstituted by transfecting chicken DF-1 (A) and human HEK-293T (B) cells and incubating at 39°C and 37°C, respectively. Luciferase activities were measured 24 hrs post-transfection. RNP complexes without PB1 served as negative controls. The polymerase activity of Anhui/13 H7N9 was set at 100% and percent relative luciferase units (% RLU) was calculated. The data shown are representative of two independent experiments. Ordinary One-way ANOVA was carried out by comparison with Anhui/13 H7N9. * denotes P-value <0.05; ** denotes P value <0.005; *** denotes P-value <0.001; **** denotes P-value <<0.0001.

In human cells (HEK-293T), the vRNP of Anhui/13 showed statistically significant higher polymerase activity compared to the vRNPs of Pakistan, Bangladesh and Vietnam (Figure 6B). Among the H9N2 vRNP complexes, Pakistan exhibited the highest polymerase activity, followed by Vietnam, with the vRNP of Bangladesh exhibiting the lowest polymerase activity. The PA alone, or in combination with PB2 on Anhui/13 H7N9 background, also showed reduced polymerase activity in human cells. This observation suggested that the vRNP of Anhui/13 is more active in human cells, followed by Pakistan and Vietnam, while Bangladesh is relatively less dynamic in human cells.

## Discussion

The China-origin H7N9 LPAIV emerged in China in 2013 and continued to reassort with local enzootic H9N2 viruses (2, 28). We modelled potential outcomes of reassortment events between H7N9 and the H9N2 viruses enzootic in poultry in the wider Asian geographical regions. The G1 lineage is prevalent in birds in the Indian subcontinent, the Middle East and Africa, whereas BJ94-like viruses are enzootic in China and neighbouring countries. To test the propensity of H7N9 acquiring internal genes from other co-circulating H9N2 viruses, three 2:6 H7N9 RG viruses were generated which contained external HA and NA genes of the China-origin H7N9 and internal gene segments from three different enzootic H9N2 AIVs. Chickens infected with these three reassortants showed clear differences in their replication kinetics and transmissibility. Both the Bangladesh_2:6_ and Pakistan_2:6_ contained G1 lineage internal gene cassettes, but differed in their transmission efficiency, with the former clearly failing to transmit (i.e. 0% transmission to contacts) while the latter demonstrated 67% transmission efficiency to contacts. While original H9N2 Pakistan virus transmitted efficiently (100%) and rapidly to contact chickens as observed in previous studies (36–38), coupling of its internal genes with the HA and NA from Anhui/13 reduced the transmission efficiency, despite a higher polymerase activity in chicken cells for the Pakistan vRNP complex compared to that of Anhui/13 (34). It is likely that internal gene cassettes from H9N2 viruses akin to the Bangaldesh_2:6_ and Pakistan_2:6_ would be unable to establish sustained circulation in chickens, and would therefore not contribute to any prolonged changes in H7N9 epidemiology. The reduced infectivity and transmissibility of the 2:6 H7N9 viruses could be due to incompatibilities between the surface glycoproteins and the internal genes, either through protein or RNA interactions affecting packaging signals. Indeed, packaging signal incompatibilities have been observed to hinder the fitness of reassortment viruses generated between China origin H7N9 and human H3N2 viruses (39) and clearly plays an important part in the emergence of novel genotypes through reassortment (40). By contrast to the two 2:6 H7N9 reassortants with G1 H9N2 internal genes (Pakistan_2:6_, Bangaldesh_2:6_), the Vietnam_2:6_ most closely resembled the ability of Anhui/13 to both shed (notably from the cloacal tract) and transmit successfully at 100% efficiency. The close phylogenetic similarity of internal genes originating from Anhui/13 (G57-origin) and the Vietnam H9N2 compared to those of the other two G1 lineage H9N2 AIVs may at least partly explain this observation (Supplementary Figure 1).

To investigate the contribution of internal gene segments on viral fitness, we performed three *in vivo* co-infection experiments in chickens. We have previously generated a large range of reassortant viruses *in vivo* between Anhui/13 and the Pakistan H9N2 AIV, where the progeny included H9N2 and a novel H9N9 virus subtype, despite 1000-fold lower titres of Pakistan virus in the initial mixed inoculum (34). The current study featured co-infections of the 2:6 H7N9 reassortants with the unaltered prototype H7N9 (Anhui/13), where any reassorted viable progeny viruses would be constrained to possess the Anhui/13 HA and NA external genes (i.e. retaining the H7N9 subtype), but with the potential to include novel genotypes involving the internal gene cassettes of G57-origin (Anhui/13), G1-like or BJ94-like lineage H9N2 AIVs. Importantly, the *in vivo* dynamics of the range of potential new reassortants was assessed through to the R2 transmission stage, at which point the shedding of exclusively unaltered Anhui/13 was detected following the three initial co-infections. We therefore demonstrated the favourable combination of the Anhui/13 external genes along with all six of its G57-origin internal genes, which at the R2 stage had successfully outcompeted any new genotypes *in vivo*. The identification of novel influenza A virus threats to animal and human health may be at least partly dependent on understanding the replicative advantages conferred by internal gene cassettes such as those of G57-origin.

Despite an ultimate selective advantage for Anhui/13 internal genes, a range of H9N2-origin internal genes were identified as transient H7N9 viable genotypes during shedding at the D0 and R1 stages. The five novel H7N9 genotypes all possessed at least one of the three polymerase genes of either Vietnam or Bangladesh origin, but there was no detectable transmission of any of the Pakistan- origin internal genes (Figure 3 and Table 1). The previously observed viral progeny from the Anhui/13 and H9N2 Pakistan co-infections frequently included the H9 HA gene (34), which again suggested that gene incompatibilities existed between the H7 HA gene and Pakistan-origin internal genes in the current study. The relative abundance and range of transient novel genotypes was relatively low for all co-infections, despite the administration of equal infectious levels of both viruses in the three inocula. However, the level of reassortment was greatest for the BJ94-like virus (Vietnam_2:6_) co-infection, where a higher abundance of Vietnam-origin gene segments was also detected in the cloacal swab samples. This observation was consistent with the increased cloacal shedding observed in the single-virus infections with Vietnam_2:6_. We identified the PB2, NP and M gene segments from the BJ94-like lineage virus to be abundantly shed from the oropharyngeal cavity of the D0 chickens at the later time point. Interestingly, the PB2, NP and M gene segments have also been identified as the key virulence genes for Anhui/13 in mice (41).

In the case of the co-infections which featured the H9N2 G1-like lineage internal genes (Pakistan_2:6_ and Bangladesh_2:6_), the G1-origin NP was absent in all D0 oropharyngeal samples, even as early as 1 day post infection, indicating that the Anhui/13 NP is immediately selected in preference to G1-like NP gene segments. Interestingly, in the field, sequence analysis indicated that H7N9 has inherited NP genes from co-circulating H7N9 viruses instead of H9N2 viruses, despite frequent co-circulation of H9N2 viruses in poultry in China (42). Moreover, mutational analysis indicated divergent evolutionary pathways between NP genes from H7N9 and H9N2 viruses (42). Furthermore, the 2014 human H5N6 virus isolate first reported in the Sichuan Province of China contained an NP gene segment with 99% similarity to that in contemporary H7N9 and H10N8 viruses (42). Together, these observations support our findings that the NP gene segment from Anhui/13 is more optimal in chickens than the NP of the G1-lineage viruses and may readily replace other NP segments during reassortment events in the field. However, no significant increase in polymerase activity was associated with Anhui/13-origin NP in chicken cells in the current study (Supplementary Figure 7), or indeed previously (34). These observations indicate that the role of Anhui/13-NP in reassortment merits further investigation.

Multi-step replication kinetics in chicken cells showed that the reassortants Vietnam_2:6_ and H7N9-R1-V31 replicated at significantly higher titers compared to Anhui/13. This increased replication is partially attributed to the higher polymerase activity of Vietnam-origin internal genes, and implied that H7N9 reassortment with G57-lineage H9N2 viruses may occur in chickens to produce viable viruses possessing an increased replicative fitness in this host. The multi-step replication kinetics in human cells showed significantly increased replication associated with Anhui/13 H7N9 compared to most of the reassortant viruses, which also correlated with the higher polymerase activity of Anhui/13 in human cells. Moreover, *in silico* analysis of the cumulative number of humanising polymorphisms present in every potential reassortment virus used in this study revealed that the wt Anhui/13 still had the greatest number, and thus all other potential reassortants posed a lower risk to human health. However, the two reassortant viruses H7N9-R1-V29 and H7N9-R1-V31 (containing Bangladesh- and Vietnam-origin H9N2 internal genes respectively) showed increased and comparable replication to Anhui/13 in human cells respectively. Thus, in the case of both G57-like and G1-like lineage H9N2 viruses reassorting with H7N9 viruses, there is a possible outcome which includes the emergence of novel viruses with a potentially increased threat to human health. However, the human threat of these viruses was only assessed *in vitro*, and investigation *in vivo* would be required to appreciate the true public health risk.

Among the five reassortant viruses recovered from three R1 chickens, three viruses originated from the Bangladesh_2:6_ while two viruses originated from Vietnam_2:6_ co-infected groups. Interestingly, two of these R1 chickens harboured multiple genotypes in the swab samples. In addition, all three chickens also shed a genetic constellation which differed between the oropharyngeal and cloacal cavity from the same bird. This observation was surprising because transmission is often considered to represent a ‘bottle neck’ to viral (including influenza A virus) diversity, with only limited virus genotypes being transmitted to successfully establish infection in the new host due to the triggering of local immune responses (43, 44). However, the effect of the bottle neck may vary, depending on species and transmission route (45). In avian hosts, the replication and dissemination of progeny AIV occurs via both the respiratory and enteric tracts, suggesting that two anatomically separate bottle necks may function simultaneously in infected birds.

Several studies have investigated the consequences of H7N9 and H9N2 virus reassortment by generating a panel of viruses by reverse genetics and focused on potential zoonotic outcomes using mouse models of infection (41, 46). We differed by investigating the consequences of mixed infections of H7N9 subtypes in chickens, arguably the species where novel reassortant viruses are most likely to be generated and maintained, with subsequent potential spill-over to humans. In addition to our previous study (34), another investigation has described *in vivo* reassortment in chickens between H7N9 and H9N2 viruses (47). The authors adopted a similar approach to our previous study (34) by using whole *wt* AIVs, with the corresponding *wt* HA and NA gene segments (i.e. H7, H9, N9 and N2) (47). They reported a greater range of reassortment between two China-origin H7N9 viruses and G57-like, G44-like and G1-like H9N2 viruses in comparison to our current study. However, both studies concurred in that emerging reassortants became rapidly outcompeted by the H7N9 *wt* variants during chicken transmission (47). The studies differed concerning the detail of reduced transmissibility of reassortant viruses, whereby we previously showed that H7N9 *wt* co-infection with a more contemporary G1-like virus generated H9N2 and H9N9 reassortants which transmitted efficiently in chickens (34). In this study, we focused on the role of the internal genes, removing the confounder of the HA and NA gene segments, and compared the H9N2 internal gene cassettes in isolation. Importantly, our co-infection approach provided opportunity for replicative competition between internal genetic segments. In contrast to W. Su et al. (47), we observed that H7N9 viruses possessing G1-like and BJ94-like are capable of efficient transmission in chickens, but were rapidly outcompeted when competition was provided by H7N9 co-infection. To our knowledge, we have reported the first description of co-infection and transmission outcomes resulting from different internal gene constellations from H7N9 and several different H9N2 viruses.

While widespread vaccination of poultry in China has largely controlled human cases of H7N9 (42), this having been implemented following the emergence of an H7N9 HPAIV derivative (48). H7N9 has been sporadically detected in poultry and the environment in farms and live bird markets across different provinces of China (Fujian, Guangdong and Henan) between January 2020 and October 2021 (24), suggesting continued circulation in Chinese poultry. Our data suggested that while the Anhui/13 internal gene cassette is optimal for transmission of the China-origin H7N9, reassortment with other H9N2 viruses can occur in chickens to produce viable viruses capable of transmission. In addition, these viruses may have altered biological properties which may elevate their zoonotic threat.

## Materials and Methods

### Ethics Statement

All animal studies and procedures were carried out in strict accordance with European and United Kingdom Home Office legislation. As part of this process, the *in ovo* and *in vivo* work was subject to scrutiny and approval by the Animal Welfare Ethical Review Board (AWERB) at The Pirbright Institute (TPI) and the Animal and Plant Health Agency (APHA), Weybridge, respectively. Any infected poultry which began to display severe clinical signs were euthanised and were recorded as a mortality. UK regulations categorise the H7N9 LPAIV as a SAPO 4 and ACDP Hazard Group 3 pathogen because it is a notifiable animal disease agent and presents a zoonotic risk. All laboratory and infectious work with H7N9 specimens, including infected poultry and eggs, was done in licenced ACDP3/SAPO4 laboratories of TPI or APHA.

### Influenza A viruses prepared by reverse genetics and their propagation

The nucleotide sequences of the different gene segments of H7N9 and all H9N2 viruses were retrieved from publicly accessible databases, namely the Global Initiative on Sharing All Influenza Data (GISAID https://www.gisaid.org/) or the National Center for Biotechnology Information (NCBI https://www.ncbi.nlm.nih.gov/). Gene segments were synthesised by Geneart^TM^ (Thermo-Fisher Scientific) and subcloned into the pHW2000 vector by restriction enzyme dependent (49), or restriction enzyme and ligation independent (50, 51), cloning techniques. The four influenza viruses used in the study included prototype China-origin H7N9 A/Anhui/1/13 (abbreviated to Anhui/13, GISAID accession no. EPI_ISL_138739) and three H9N2 viruses which supplied the six internal gene segments for 2:6 reassortment with the haemagglutinin (HA) and neuraminidase (NA) of Anhui/13, namely: A/chicken/Pakistan/UDL-01/2008 (NCBI Accession number; CY038455), A/Environment/Bangladesh/26218/2015 (NCBI Accession number; KY635657.1), and A/chicken/Vietnam/H7F-14-BN4-315/2015 (GISAID Accession number; EPI_ISL_327772). The 2:6 reassorted viruses were rescued by reverse genetics (36, 52), and propagated in 10-day-old specific pathogen free (SPF) embryonated chicken eggs at 37 °C for 72 hours. The Madin Darby canine kidney (MDCK) and human embryonic kidney (HEK) 293T cells (ATCC) were maintained with Dulbecco’s Modified Eagle’s medium (DMEM) (Sigma), supplemented with 10% fetal calf sera (FCS) (Sigma), 100U/mL penicillin, and 100μg/mL streptomycin (Gibco) at 37°C under 5% CO_2_ (v/v).

### 50% egg infectious dose 50 (EID_50_) determination

Egg infectious dose of the viruses was calculated by making ten-fold serial dilutions of the viruses. 100μl of each dilution was inoculated in a group of 6 eggs and eggs were incubated at 37°C for 72 hours (53). Allantoic fluid was harvested from all the inoculated eggs and tested for the presence or absence of virus by the haemagglutination assay (53). EID_50_ titre was calculated by the method described by Reed and Muench (54). The viruses were aliquoted and stored at −80°C till further use.

### AIV infections of chickens

The specific pathogen free (SPF) chickens of Rhode Island Red variety (procured from the National Avian Research Facility (NARF), Roslin Institute, Edinburgh, UK) at three weeks of age were used in the experiment. For the single-virus infections, three groups of six chickens were infected intranasally (i.n.) with 200μl of inoculum (applied across both nares of each directly-infected (D0, with “D” denoting “donor”) chickens containing 1×10^8^ EID_50_ of the three 2:6 reassortant viruses (i.e. with Pakistan-, Bangladesh- or Vietnam-origin internal genes), and a fourth group of six D0 chickens were similarly infected with 1×10^8^ EID_50_ of Anhui/13. Contact chickens (n=18), referred to as ‘R1’ (“R” denotes “recipient”), were introduced for co-housing with D0 chickens at 1-day post-infection (dpi). For the co-infections, D0 chickens (n=6/group) were co-infected i.n. with 200μl of inoculum containing a mixture of the 1X10^8^ EID_50_ of Anhui/13 and 1×10^8^ EID_50_ of the reassortant 2:6 H7N9 viruses. R1 chickens (n=6/group) were introduced for co-housing at 1 dpi. When transmission to 50% of the R1 chickens had occurred, all D0 chickens were removed from cohousing by culling, and the second group of contact chickens, referred to as ‘R2’, were added for cohousing at 4 or 5 dpi. For single-virus and co-infected chickens, oropharyngeal and cloacal swabs were collected daily in Virus Transport Medium (VTM) (34) from all the directly infected chickens and contact chickens until 14 dpi. Chickens were monitored for clinical signs up to three times daily. Any surviving chickens were humanely euthanized by overdose of pentobarbitone at 14 dpi, with blood collected by terminal heart bleeds. Time points relating to the R1 and R2 contact chickens may at times be referred to as days post-contact (dpc), as appropriate.

### Serology

Sera derived from clotted blood were heat inactivated at 56 °C for 30 minutes. Seroconversion in the infected chickens was identified via the haemagglutination inhibition (HI) test (53), using the homologous inactivated Anhui/13 virus as antigen titrated to 4 haemagglutination units.

### Avian influenza virus Screening RT-qPCR

RNA was extracted from VTM swab fluids by robotic extraction using the RNeasy RNA extraction kit (Qiagen), as described previously (55, 56). For initial screening purposes, all extracted RNA specimens were tested by the M-gene RT-qPCR using the primers and probe, as described previously (57, 58). A ten-fold dilution series of titrated H7N9 RNA was used to calculate viral titres were which are displayed as relative equivalency units (REUs) (34).

### H7N9/H9N2 gene specific RT-qPCR

Segment specific primer and probes for Pakistan, Bangladesh, and Vietnam viruses were designed to distinguish between the H7N9- and H9N2-origin internal gene segments, as described previously (34, 55). Briefly, each segment-specific assay consisted of a pair of conserved primers which bind to identical sequences which are shared within each segment of Anhui/13 and an H9N2 virus. These conserved primers generated amplicons which were distinguished as regards their viral origin by using probes which were targeted to strain-specific nucleotide sequences within the amplicons. Each probe was 5’-labelled with a different fluorophore, namely FAM and HEX for Anhui/13 and H9N2-origin segments, respectively (Supplementary Table 1). All assays were adapted to the One-step RT-PCR Kit (Qiagen) reaction chemistry, as described previously and used the same thermocycling conditions (34). Each segment-specific RT-PCR was compared for its analytical sensitivity to demonstrate equivalent performance of the assays (Supplementary Figure 6). RNA from oropharyngeal and cloacal samples was analysed by both RT-qPCR assays to calculate the relative percentage abundance (origins) of each gene segment.

### Plaque purification

MDCKs were infected with a ten-fold serial dilution of a swab suspension made in 500 μl of DMEM and incubated for 1 hour at 37 °C. The infection media were removed and 2 ml of overlay media with 2 % agar added, inverted, and incubated at 37 °C with 5% CO_2_ for 4 days, or until plaques are formed. Plaque-purification was performed in quadruplicate, in six-well plates. Discrete plaques were identified and harvested into 200 μl of serum free media using a pipette tip pushed through the layer of agar until contact with the surface of the plate was made. RNA was extracted from the media plaque suspensions by robotic methods described previously (55, 56).

### Growth kinetics of reassortant viruses

Growth kinetics of the reassortant viruses was assessed in DF-1 cells (chicken embryo fibroblast), A549 (human alveolar basal epithelial) and Calu-3 (human lung) cells. Growth kinetics was performed using 12-well plates in quadruplicate. Cells at 90% confluency were infected with 0.001 multiplicity of infection (MOI) of respective viruses in DMEM containing antibiotics and 0.3% BSA. The cell supernatant was harvested after 24, 48 and 72 hrs post-infection and titrated by TCID_50_ in MDCK cells (54).

### Minireplicon assay

The polymerase activity of different vRNP combinations was assessed *in vitro* by plasmid-based reporter gene expression in 24-well plates (59). Chicken DF-1 cells and human HEK-293T were transfected with expression plasmids for different RNP combinations prepared from the Anhui/13, Pakistan, Bangladesh and Vietnam progenitor viruses by using Lipofectamine 2000 (Invitrogen), as per the manufacturer’s instructions. 80 ng of pCAGGS plasmid encoding PB2 and PB1, 40 ng of pCAGGS plasmid encoding PA and 160 ng of pCAGGS plasmid encoding NP were co-transfected with 40 ng of a Renilla Luciferase pCAGGS expression plasmid and 80 ng of a pCk-PolI-Firefly plasmid. The cells were incubated for 24 hrs at 37 °C (for HEK-293T) or 39 °C (DF-1) and luciferase activity was measured by using the Dual-Glo luciferase assay system (Promega). The polymerase activity was calculated by normalising firefly luciferase activity relative to the Renilla luciferase activity.

## Funding

This research was funded by BBSRC grant numbers (BB/N002571/1, BB/S013792/1, BBS/E/I/00007034, BBS/E/I/00007035), Zoonoses and Emerging Livestock systems (ZELS) (BB/L018853/1 and BB/S013792/1), the GCRF One Health Poultry Hub (BB/S011269/1), UK-China-Philippines-Thailand Swine and Poultry Research Initiative (BB/R012679/1), plus Defra (UK, including the Devolved Administrations of Scotland and Wales) for the work programmes SE2211 and SE2213. The funders had no role in study design, data collection and analysis, decision to publish, or preparation of the manuscript.

Author contributions were as follows. Conceptualization -SB and MI. Formal Analysis –JJ and SB. Funding Acquisition–MI. Investigation–JJ, SB, SKW, HMTKK, JRS, PC, JES, SM, BCM, MJS, SMB and MI, Methodology– JJ, SB, SKW, HTMTKK, MJS and MI. Resources–SB, JJ, MJS, SMB and MI. Supervision–MI. Visualisation–JJ and SB. Writing –Original Draft Preparation–JJ and SB. Writing –Review and Editing–JJ, SB, MJS, HMTKK, PC and MI.

## Competing interests

The authors declare they have no conflict of interest.

## Acknowledgements

We would also like to thank Caroline Warren, Alexander MP Byrne and Saumya Thomas from Animal and Plant Health Agency for help processing samples. We would like to thank Holly Shelton and Dagmara Bialy for their support for the ACDP3/SAPO4 work in the Avian West Wing of The Pirbright Institute.

---

**Supplementary Figure 1. Phylogenetic analysis of Haemagglutinin (HA) genes of H9N2 AIVs and a schematic of the viruses used in this study. (A)** Phylogenetic relationship of the HA genes of the H9N2 AIVs which were used to generate the 2:6 reassortant H7N9 viruses by reverse genetics (RG). **(B)** The identify of the viruses is shown in different colours; red, Pakistan; green, Vietnam; purple, Bangladesh; blue, Anhui/13 or its ancestral H9N2 virus (G57). Phylogenetic tree was generated by maximal-likelihood analysis using H9Nx sequences downloaded from GISAID Epiflu database.

**Supplementary Figure 2. Phylogenic analysis of the internal genes of the H9N2 AIVs which provided internal genes for this study.** A maximal likelihood phylogenetic tree of all genes of the viruses including several reference viruses representative of the different dominant clades. The phylogenetic location of the viruses used in the study is shown in different colours; blue, Anhui; red, Pakistan; green, Vietnam; purple, Bangladesh. * HA and NA gene segments shown were from the closest related H9N2 virus which naturally donated its internal gene cassette to produce the prototype H7N9 Anhui/13.

**Supplementary Figure 3. Viral shedding and transmission of Anhui/13 or reassorted 2:6 H7N9 viruses possessing different H9N2 internal genes in individual chickens.** Influenza virus titres from oropharyngeal swabs of chickens (n=6 per group) infected with 1×10^8^ EID_50_ of Anhui/13 **(A)** or 2:6 H7N9 viruses possessing the internal genes from either Vietnam **(B)**, Pakistan **(C)** or Bangladesh **(D)** H9N2 viruses. Oropharyngeal shedding titres from chickens placed in-contact with infected groups at 1 dpi **(E-H)**. Viral shedding displayed as relative equivalency units (REUs) based on M-gene RT-qPCR derived from a standard curve (dilution series) of viral RNA from titrated Anhui/13. Each coloured line represents shedding from an individual chicken. Dotted horizontal line indicates the positive cut-off at a Ct value of 36.

**Supplementary Figure 4. H7N9 seroconversion of chickens infected with Anhui/13 or 2:6 H7N9 viruses possessing different H9N2 internal genes.** Sera from D0 and R1 chickens infected with Anhui/13 or each of the different 2:6 H7N9 viruses (n=6 per group), collected at 14 dpi and tested by the HI assay using the H7N9 Anhui/13 antigen. Individual titres with geometric mean +/− SD are shown. Dotted line represents the HI positive cut-off threshold titre at 1/16.

**Supplementary Figure 5. Seroconversion of chickens placed in-contact with chickens co-infected with Anhui/13 and an individual 2:6 H7N9 virus possessing different internal genes.** Seroconversion from R1 and R2 chickens was measured in a transmission chain from chickens co-infected with Anhui/13 or each of the different 2:6 H7N9 viruses, with six chickens per group. Sera were collected at 14 dpi and tested HI using H7N9 Anhui/13 antigen. Individual titres with geometric mean +/− SD are shown. Dotted line represents the HI positive cut-off threshold titre at 1/16.

**Supplementary Figure 6. Comparative sensitivity of the gene segment specific RT-qPCRs.** Influenza segment and virus specific RT-qPCRs developed for Anhui/13 (HEX fluorescence) and the 2:6 reassorted H7N9 viruses with internal genes from Vietnam, Pakistan or Bangladesh H9N2 AIVs (FAM florescence). The pairs of assays were tested for equivalence against 10-fold serial dilutions of viral RNA extracted from each virus, starting at 1×10^6^ EID_50_, and are shown for (A) the segment-specific RT-qPCRs which distinguish the viral origin, and for the (B) generic M-gene RT-qPCR which served to quantify the total viral shedding during the *in vivo* experiments. Ct values are plotted against virus titre, lines of best fit are displayed. Dotted lines represent the positive cut-off at Ct 36.

**Supplementary Figure 7. Minireplicon assays of the ribonucleoprotein (RNP) complexes of H7N9 Anhui/13 and H9N2 UDL/08, H9N2 Vietnam/315, H9N2 Bangladesh/26218 and their additional RNP combinations.** The RNP complexes were reconstituted by transfecting chicken DF-1 (A) and human HEK-293T (B) cells and incubating at 39°C and 37°C, respectively. Luciferase activities were measured 24 hrs post-transfection. RNP complexes without PB1 served as negative control. The polymerase activity of Anhui/13 H7N9 was set at 100% and percent relative luciferase units (% RLU) was calculated. The data shown are a representative of two independent experiments. Ordinary One-way ANOVA was carried out by comparison with Anhui/13 H7N9. * denotes P-value <0.05; ** denotes P value <0.005; *** denotes P-value <0.001; **** denotes P-value <0.0001.

**Supplementary Figure 8. Multi-step replication kinetics of Anhui/13 H7N9, 2:6 H7N9 viruses and the reassortant viruses in human Calu-3 cells.** Virus replication of 2:6 H7N9 viruses containing the HA and NA from H7N9 and the remaining internal gene cassettes from either Pakistan, Vietnam or Bangladesh (A-C) H9N2 viruses or novel reassortant viruses recovered by plaque purification from R1 chicken swabs (D-H), see Table 1. Cells were infected at a MOI of 0.01 with each virus and cell supernatant was harvested at 24, 48 and 72hr post-infection. Viral titres in the cell supernatant were determined by TCID50/ml. Each time point corresponds to the mean of four biological replicates with standard deviations indicated. Replication kinetics of reassortant viruses compared to parental Anhui/13 H7N9 virus is shown in all panels A-H. The genotype of each reassortant virus is shown as a combination of black, red, green and violet colours. Two-way ANOVA with multiple analysis was performed comparing every group to Anhui; * indicates P-value = 0.0237; ** indicates P-value < 0.005, **** indicates P-value <0.0001.

